# Validation of a low-cost, carbon dioxide-based cryoablation system for percutaneous tumor ablation

**DOI:** 10.1101/454413

**Authors:** Bailey Surtees, Sean Young, Yixin Hu, Guannan Wang, Evelyn McChesney, Grace Kuroki, Pascal Acree, Serena Thomas, Tara Blair, Shivam Rastogi, Dara L. Kraitchman, Clifford Weiss, Saraswati Sukumar, Susan C. Harvey, Nicholas J Durr

## Abstract

Breast cancer rates are rising in low- and middle-income countries (LMICs), yet there is a lack of accessible and cost-effective treatment. As a result, the cancer burden and death rates are highest in LMICs. In an effort to meet this need, our work presents the design and feasibility of a low-cost cryoablation system using widely-available carbon dioxide as the only consumable. This system uses an 8-gauge outer-diameter needle and Joule-Thomson expansion to percutaneously necrose tissue with cryoablation. Bench top experiments characterized temperature dynamics in ultrasound gel demonstrated that isotherms greater than 2 cm were formed. Further, this system was applied to mammary tumors in an *in vivo* rat model and necrosis was verified by histopathology. Finally, freezing capacity under a large heat load was assessed with an *in vivo* porcine study, where volumes of necrosis greater than 1.5 cm in diameter confirmed by histopathology were induced in a highly perfused liver after two 7-minute freeze cycles. These results demonstrate the feasibility of a carbon-dioxide based cryoablation system for improving solid tumor treatment options in resource-constrained environments.

## Introduction

Over 8 million cancer deaths occurred worldwide in 2012, with breast cancer being the largest cause of cancer-related mortality for women with almost 500,000 reported deaths [1]. While diagnostic and treatment technologies in the developed world have advanced such that the five-year survival rate for breast cancer is almost 90% in the United States [2], the survival rate in low- and middle-income countries (LMICs) can range from 64% in Saudi Arabia to 46% in Uganda to only 12% in The Gambia [3]. These rates are typically lower in rural areas of LMICs due to inadequate treatment and long travel times to regional hospitals [4]. The current treatment pattern used in developed countries—surgery, chemotherapy, and radiation therapy—is inefficient, inaccessible, and costly in many LMICs [5,6].

Tissue ablation has several advantages over surgical treatments for practical use in LMICs, and previous work has explored the use of cryoablation for treatment of cancers including liver, lung, prostate, and breast cancer [7–12]. Cryoablation is a minimally-invasive treatment that is well tolerated with the intrinsic anesthetic properties of cold providing local anesthesia [13]. Moreover, the ice formation can easily be tracked with ultrasound for real-time treatment guidance. These features permit precise and effective cryoablation treatment with a companion ultrasound unit and only a local skin cleaning rather than the current standard of general anesthesia and an operating theater for surgery [9,14]. Importantly, circumventing the requirement for a sterile operating room would enable treatments to be performed at local clinics, which are more accessible to patients as they are more abundant in the rural regions. Further, by virtue of being minimally invasive, cryoablation is known to reduce pain, bleeding, and recovery time when compared with surgical procedures [15].

Cryoablation kills breast cancer cells through the formation of intracellular ice crystals, which begin forming at temperatures below −20 °C; however, temperatures below −40 °C are optimal for cryoablation due to the certainty of ice crystal formation [16]. To reach these cold temperatures, argon gas is commonly stored at high-pressure and is then allowed to expand through a pinhole-like opening to reach atmospheric pressure. The gas undergoes rapid expansion and, due to the Joule-Thomson (JT) effect, rapidly decreases temperature [16]. This expansion takes place inside a cryoprobe that is percutaneously inserted into the cancerous tissue, and the temperature of the cryoprobe causes an iceball as ice crystals form in surrounding cells (Fig 1). This mechanism introduces a temperature gradient in the iceball of −40 °C at the probe tip to 0 °C at the edge of the iceball. Previous work has found that tissue held below −20 °C for over a minute creates necrosis throughout the region [16]. Treatment is cycled with two freeze cycles interrupted by a thaw cycle, which allows for both cell death via cellular dehydration as well as through the formation of intracellular ice crystals [16].

**Fig 1.**
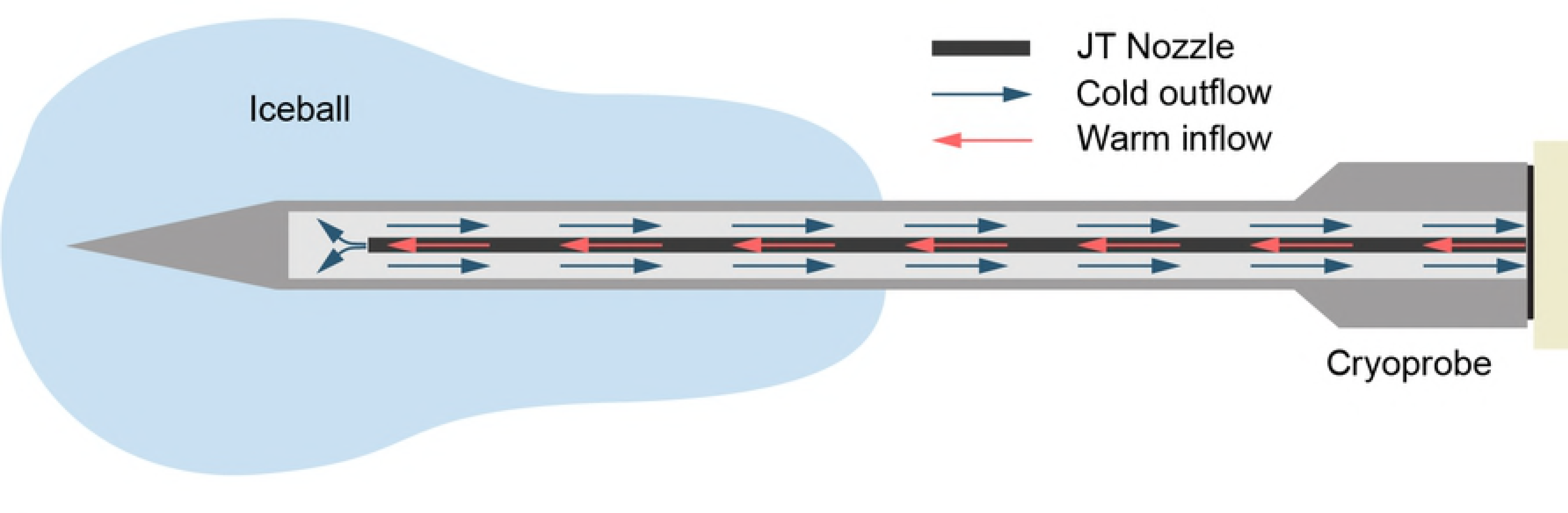
The percutaneous cryoprobe consists of a double-chamber needle. CO_2_ gas flows through the inner chamber, cools upon JT expansion at the tip, and is exhausted through the outer chamber.

Although cryoablation is promising for use in low-resource settings, most devices are not designed with cost and resource constraints in mind or for use in LMICs. Current cryoablation systems are too expensive for use in LMICs, as a single treatment can cost upwards of $10,000, with over half of the cost coming from disposable, single-use parts [17]. Furthermore, while argon gas is optimal for cryoablation, as it can reach temperatures as low as its boiling point of −185.8 °C [16], it is not available in LMICs.

Alternatively, carbon dioxide (CO_2_) is available in most rural areas due to the carbonated beverage industry and via the Joule-Thomson effect can reach a temperature as low as −78.5 °C [18]. In an effort to develop a practical system for tumor treatment in LMICs, we have assessed the performance of a custom-designed cryoablation system optimized for use with CO_2_; it is the first percutaneous cryotherapy system to use only CO_2_ as a consumable. We report our results from three stages of experimental evaluation: bench top experiment in an ultrasound gel tissue phantom, *in vivo* testing on mammary tumors in a rat model with necrosis verified via histopathology, and assessment of freezing capacity under a large heat load with an *in vivo* experiment in a porcine model.Demonstrating the feasibility of cryoablation with CO_2_ will potentially allow for increased affordable and accessible breast cancer treatment globally.

## Materials and methods

### Cryoablation system design

Our custom-designed cryosystem consists of two modules: the cryoprobe and the gas-control module. The two modules are connected with nylon tubing and joined by high-pressure brass pipefittings. The gas-control module is comprised of a high-pressure fitting that directs CO_2_ gas to the JT nozzle. This tubing allows gas release from a high-pressure environment into the cryoprobe chamber, which causes the cryoprobe to rapidly cool. The cooled CO_2_ gas then leaves the probe chamber and flows through the pre-cooling chamber that surrounds the gas inflow tube leading to the JT nozzle (Fig 2). The pre-cooling process serves as a positive feedback to the cooling system by chilling the inflow gas before it reaches the JT nozzle. After leaving the cryoprobe, the gas is released into the atmosphere through an outflow tube. The gas control module is directly connected to a tank of compressed CO_2_, and is comprised of a valve to control flow to the cryoprobe and a pressure gauge that monitors the gas supply level in the tank. To begin a freeze cycle, the valve on the CO_2_ tank is opened, and the control valve is opened to allow gas to flow through the system and cooling to begin.

**Fig 2.**
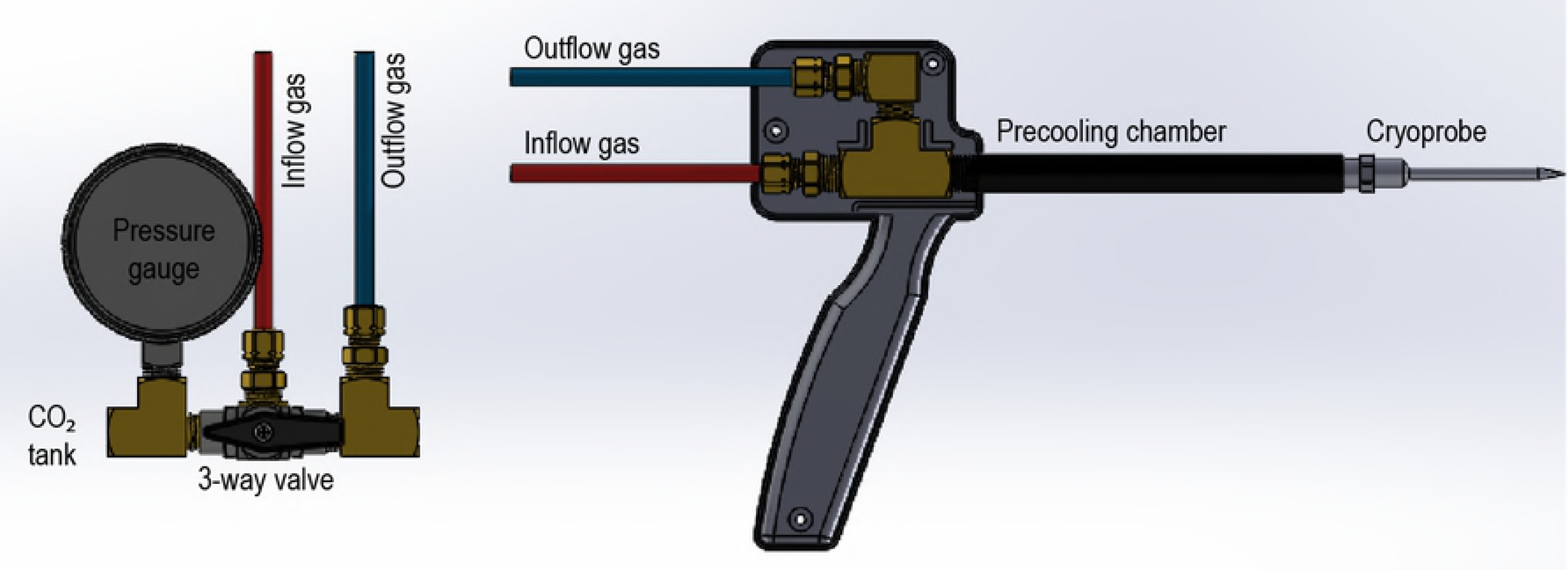
The gas control module of the cryoablation system (left) connects to a conventional compressed CO2 gas canister used in the carbonated beverage industry and regulates flow to the cryoprobe (right) during cryoablation.

### Thermodynamic profiles in tissue phantom

Initial tests were performed with the cryosystem in an ultrasound gel tissue phantom for early validation in accordance with prior studies [19]. To evaluate device performance, photographs of the iceball were taken every 30 seconds, and temperature of the cryoprobe was monitored every 0.5 seconds using a thermocouple. In total, the freezing capacity was evaluated through six trials per probe where the system completed a 5-minute freeze cycle, a 3-minute thaw cycle, and then a second 5-minute freeze cycle. For each test, the given probe was submerged in a beaker filled with 400 mL of ultrasound gel at 22.6 °C. Pictures were analyzed in ImageJ [20] to determine iceball size.

### *In vivo* cryoablation of mammary tumors

Prior to experimentation, our institutional Animal Care and Use Committee approved the rat protocol. For this portion of the study, cryoablation was performed *in vivo* in Sprague Dawley rats induced with primary mammary tumors via injections of *N*-Nitroso-*N*-methylurea (MNU). This was done to validate the efficacy of the device in necrosing breast cancer cells via histopathology, similar to previous studies [21].

Nine female Sprague Dawley rats were induced with mammary tumors by intraperitoneal injections of 50 mg/kg of MNU (analytical grade, Sigma/Oakwood Products, Inc.) at 5 weeks and 7 weeks of age. Similar to previous studies [22,23], between one and three tumors were induced per rat, each of which was treated in separate procedures. A total of ten mammary tumors in nine rats were treated by cryoablation. To determine the extent of necrosis caused by the puncture of the cryoprobe rather than the freezing, one tumor was selected at random as a no-treatment control (no freeze) in which no CO_2_ was released through the system, but all other procedural steps were followed including needle insertion and elapsed time.

All procedures were performed in the veterinary operating room aided by a trained surgeon, and each rat was anesthetized throughout the procedure. The rat was placed on a heating pad to stabilize body temperature and a rectal thermometer was used to monitor core body temperature. The tumors were surgically exposed such that each tumor was separated from the skin, keeping any large blood vessels intact. After the majority of the tumor was separated from the skin, petroleum jelly-coated gauze was placed between the tumor and the skin to act as a thermal insulator to prevent cold injury to the rat’s skin (Fig 3).

**Fig 3.**
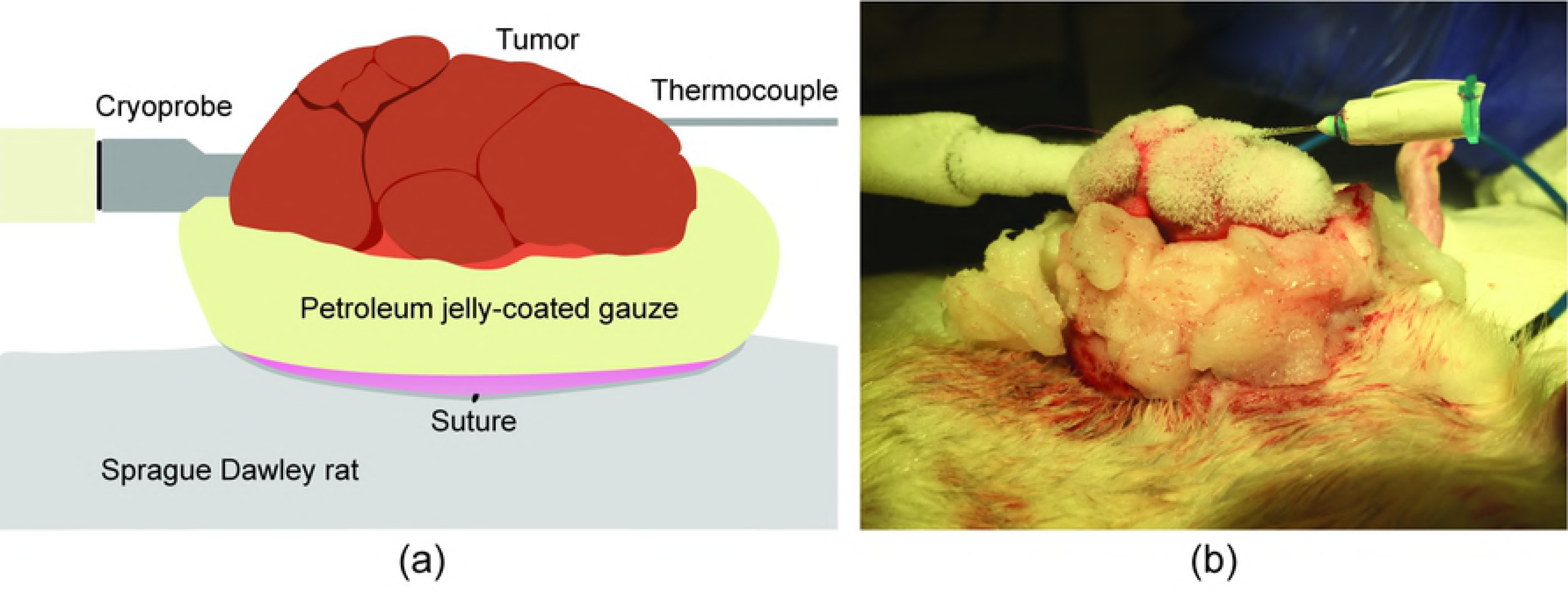
The diagram (a) depicts the treatment setup, pictured in (b), for in vivo cryoablation of mammary tumors in Sprague Dawley rats. Tumors were elevated from the body and insulated with petroleum jelly to prevent skin damage, and tumor temperature was monitored using a thermocouple during the procedure.

Prior to cryoprobe insertion, a trocar was used as a guide for the cryoprobe along the longest axis of the tumor. Then, the cryoprobe was inserted into the tumor and a thermocouple was placed 0.5 cm from the probe surface to measure tumor temperature. The control valve was then turned on and CO_2_ flowed, rapidly cooling the device and the adjacent tissue. To ensure the tumor exposure and subsequent necrosis, the first freeze cycle continued until the temperature from the thermocouple stabilized below −30 °C for at least one minute with a maximum freezing time of 7 minutes [21]. Then, the control valve was turned off for a 3-minute thaw cycle followed by the control valve being turned on for a secondary freeze cycle defined by the same temperature and time requirement as the first freeze cycle. After the second freeze, the control valve was turned off and the tumor was allowed to thaw to 20 °C followed by removal of the cryoprobe. The wounds were closed with surgical clips and the tumor remained *in situ* for a minimum of 72 hours before excision, allowing time for necrosis. The tumors were fixed in a buffered formalin fixative, and stained with hematoxylin and eosin (H&E) for examination by a pathologist to assess the extent of necrosis [21].

### *In vivo* cryoablation under heat load

Prior to experimentation, our institutional Animal Care and Use Committee approved the swine protocol. For this portion of the study, cryoablation was performed *in vivo* in the normal liver of a pig to validate the efficacy of the device under a heat load analogous to a human breast, similar to previous studies [24,25].

All procedures were performed with aseptic technique by a veterinarian and interventional radiologist. The swine was sedated and an IV catheter was introduced to administer antibiotics and anesthetic induction. The pig was intubated and general anesthesia was maintained with isoflurane anesthesia with mechanical ventilation. The swine was placed in a supine position, and a midline incision was used rather than percutaneous insertion of the probe through the skin, as the goal was to freeze only the liver tissue rather than the skin, muscle and fatty tissue anterior to the liver.

A trocar was used as a guide to enter the liver under ultrasonic guidance and then the cryoprobe was inserted. The device was positioned and CO_2_ was allowed to flow. Gauze soaked in warm saline was placed near the skin to aid in preventing skin necrosis. The iceball growth was monitored via ultrasound imaging and the timing of the freeze-thaw cycles was determined by the plateau of the iceball growth. Two 7-minute freeze cycles were performed with a 4-minute thaw cycle between the freeze cycles (Fig 4).

**Fig 4.**
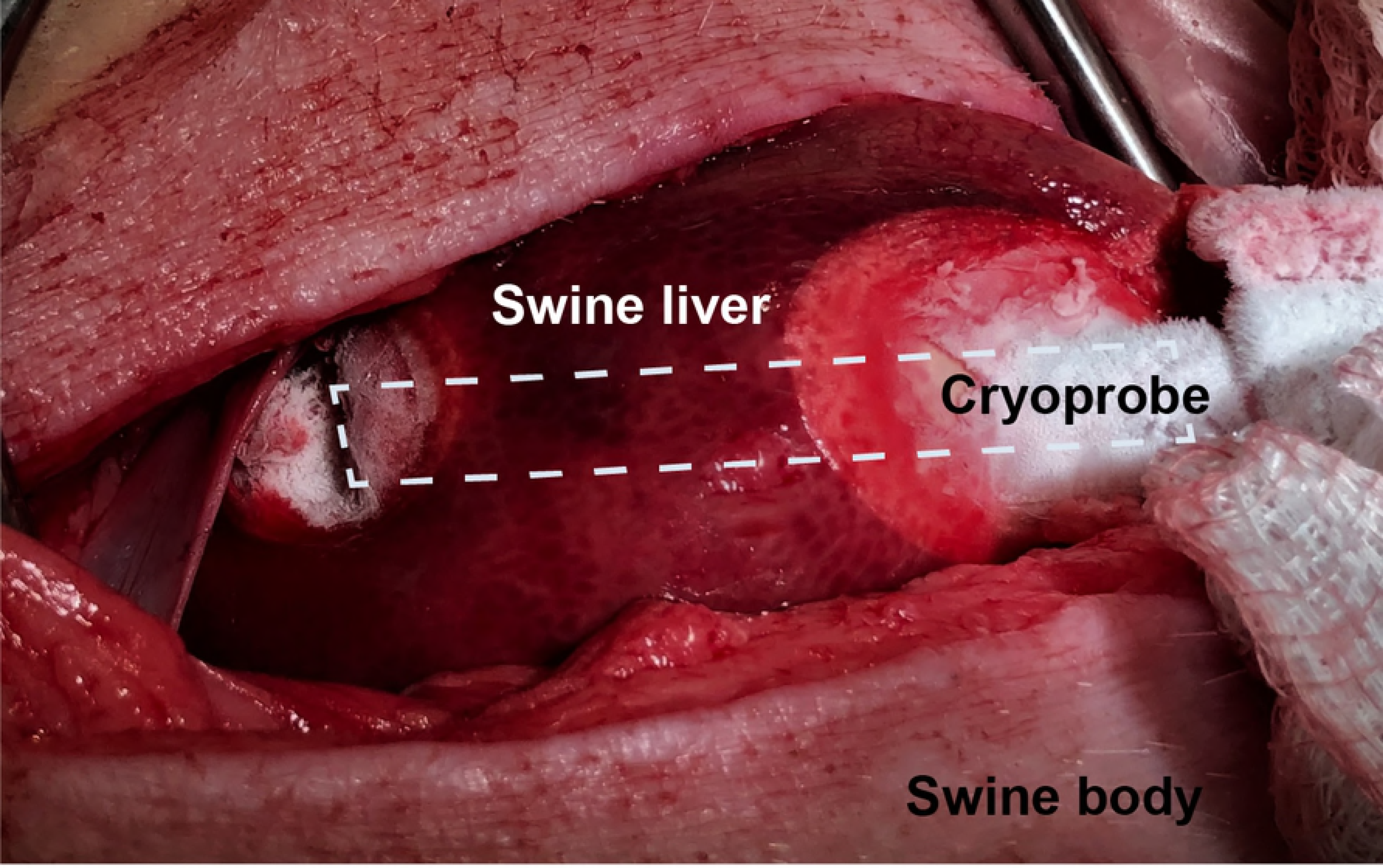
An incision was made exposing the swine liver, and the cryoprobe was inserted (outlined in white dashes). Two 7-minute freeze cycles were performed, causing an iceball to form in the liver tissue.

The midline incision was closed and the pig was recovered. The pig was humanely euthanized 48 hours after cryoablation, and the liver was excised, sectioned, and submitted for H&E staining to examine necrosis. This delay in excision and pathology was necessary as obvious signs of necrosis are not present until at least two days post-cryosurgery [21].

## Results

### Thermodynamic profiles in tissue phantom

The experiment was run on various prototypes, which guided the choice of probe for final use and further validation and are detailed below. Images taken of iceball growth over time were used to inform both final iceball size as well as isotherm size through a freeze-thaw-freeze cycle, as seen in Fig 5. Over six trials, the probe produced an average iceball diameter of 2.230 +/− 0.075 cm. Additionally, across all six trials, temperatures at the cryoprobe reached < −40 °C in the first freeze cycle, with the temperature in the second freeze cycle decreasing an additional 5°C.

**Fig 5.**
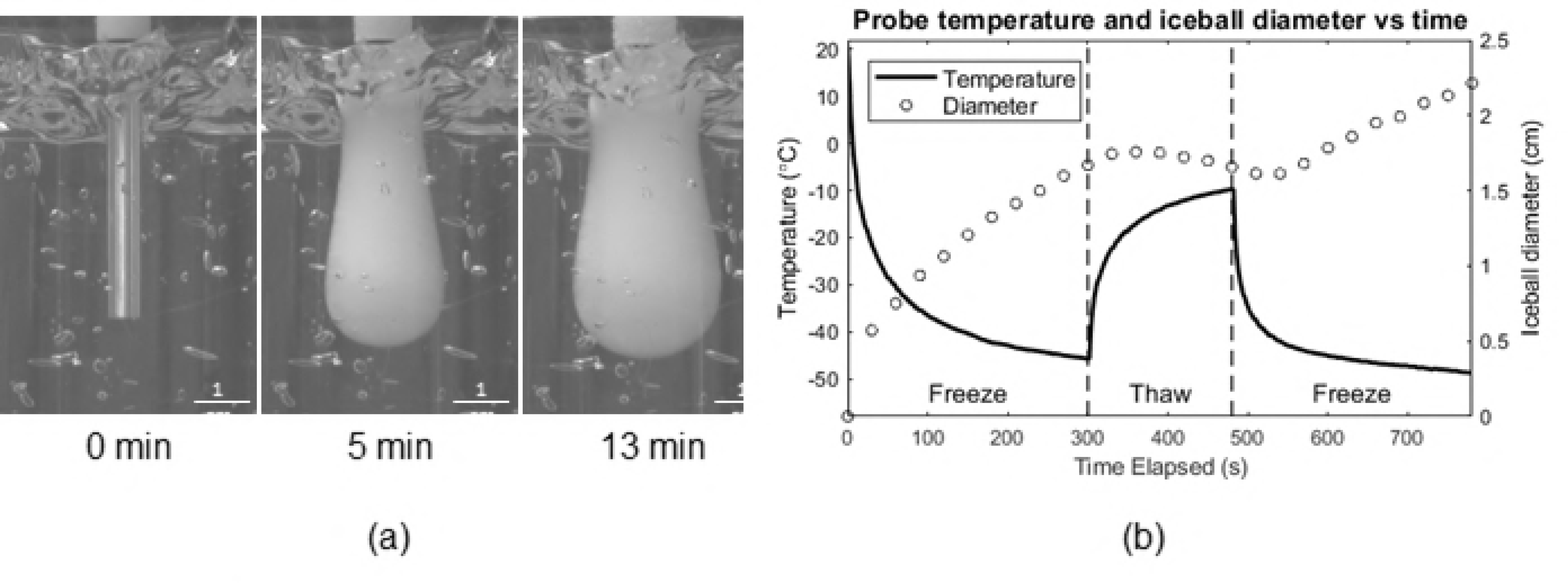
(a) Iceball growth over time as visualized in the tissue phantom, (b) plot of probe temperature at the cryoprobe tip and iceball diameter over time demonstrating temperatures below −40 °C and iceball diameters larger than 2 cm.

### *In vivo* cryoablation of mammary tumors

Two tumors were not treated using cryoablation and were instead surgically excised as it was determined that the tumors were too close to the heart, posing too great of risk to treat. Additionally, results from two trials were not included due to improper wound closure resulting in indeterminable causes of tumor necrosis. All other tumors treated in the rat experiment showed significant levels (>85%) of tumor necrosis while the tumor from the negative control trial was uniformly viable (Table 1). Tumor sizes ranged from 2.0-3.4 cm.

**Table 1.** Histological Results Categorized by Level of Necrosis.

| Region | Percent Necrosed | Tumors | Histological Characteristics |
| --- | --- | --- | --- |
| A | 100% | 5 | Tumor area completely necrotic; reactive stroma observed |
| B | ≥85% and <100% | 5 | Large area of necrotic debris, edge has a mixture of necrotic inflammatory cells, hemorrhage from cryoablation, neutrophils, degraded islands and few apoptotic cells |
| C | <85% & >0% | 0 | Moderate area of necrotic debris, mostly neutrophils, nests of viable/degrade cells |
| D | 0% | 1 Control | Viable tumor cells throughout the section |

### *In vivo* cryoablation under heat load

Tissue damage was seen in the sectioned portions of liver with defined borders between healthy and necrotic tissue (Fig 6).

**Fig 6.**
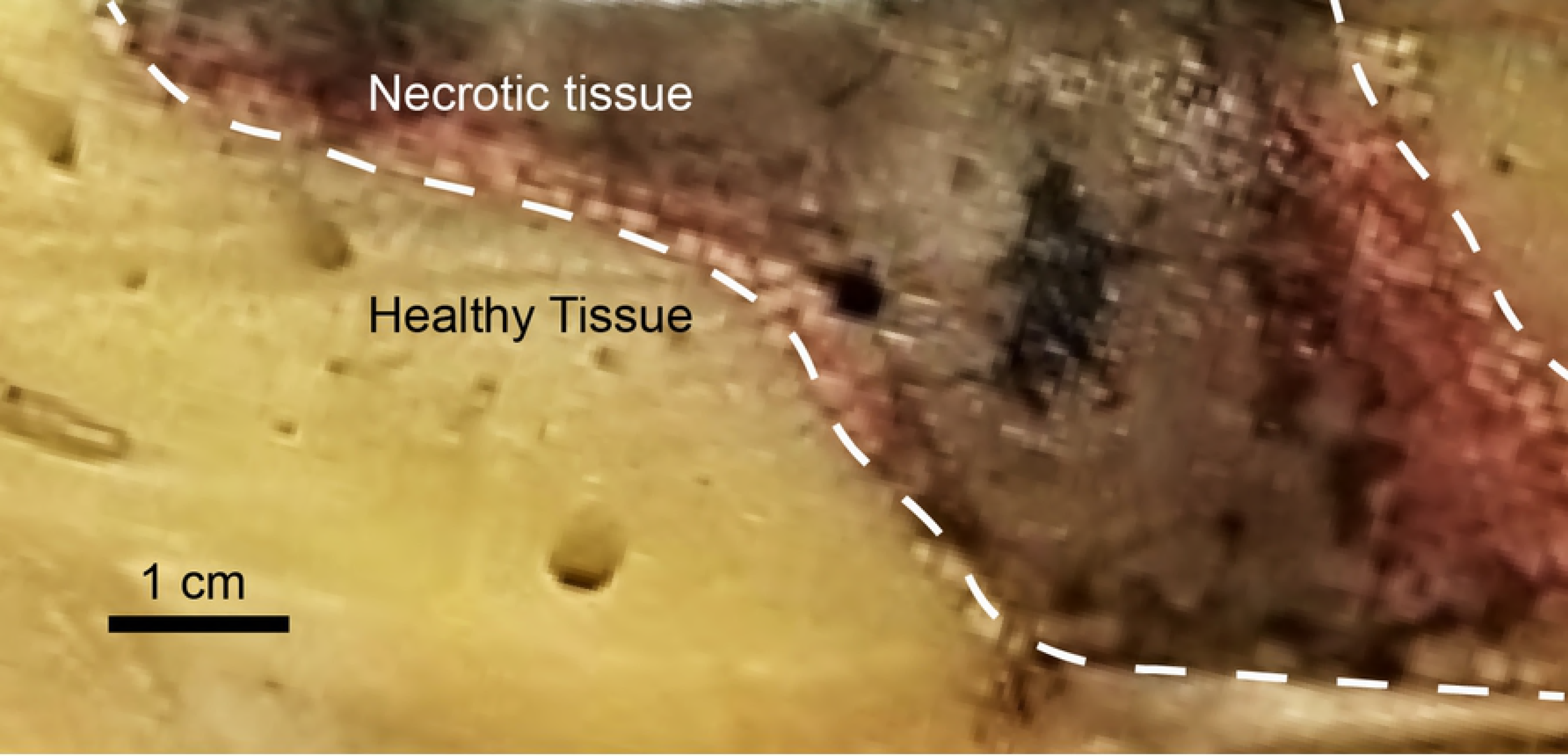
A section of the liver after perfusion and fixation where cryoablation was performed. Defined borders are seen between the dark necrotic tissue and the lighter healthy tissue.

We observed a rapid transition from necrotic tissue to relatively normal liver tissue, as is demarcated by the dashed line in Fig 7. Histopathology examination of the ablated area confirmed varied gradients of necrosis (Fig 8).

**Fig 7.**
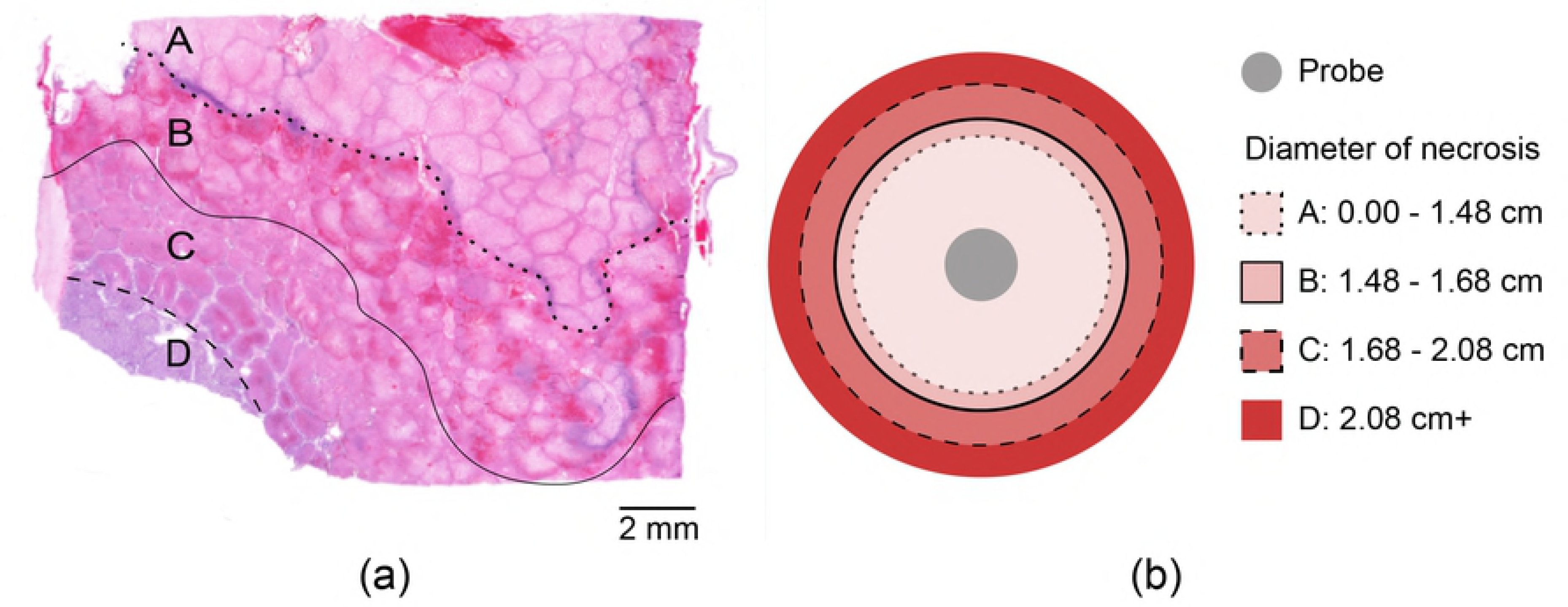
(a) Histologic analysis found a 1.48-cm region of complete necrosis (Region A), with a 0.1- to 0.2-cm wide region of complete necrosis with traces of hemorrhaging (Region B), a 0.1- to 0.4-cm margin of complete hepatocyte necrosis with intact bile ducts (Region C), and a region of non-necrotic hepatocytes beyond this (Region D) (b) Diameters of necrotic regions are depicted in a circular cross section.

**Fig 8.**
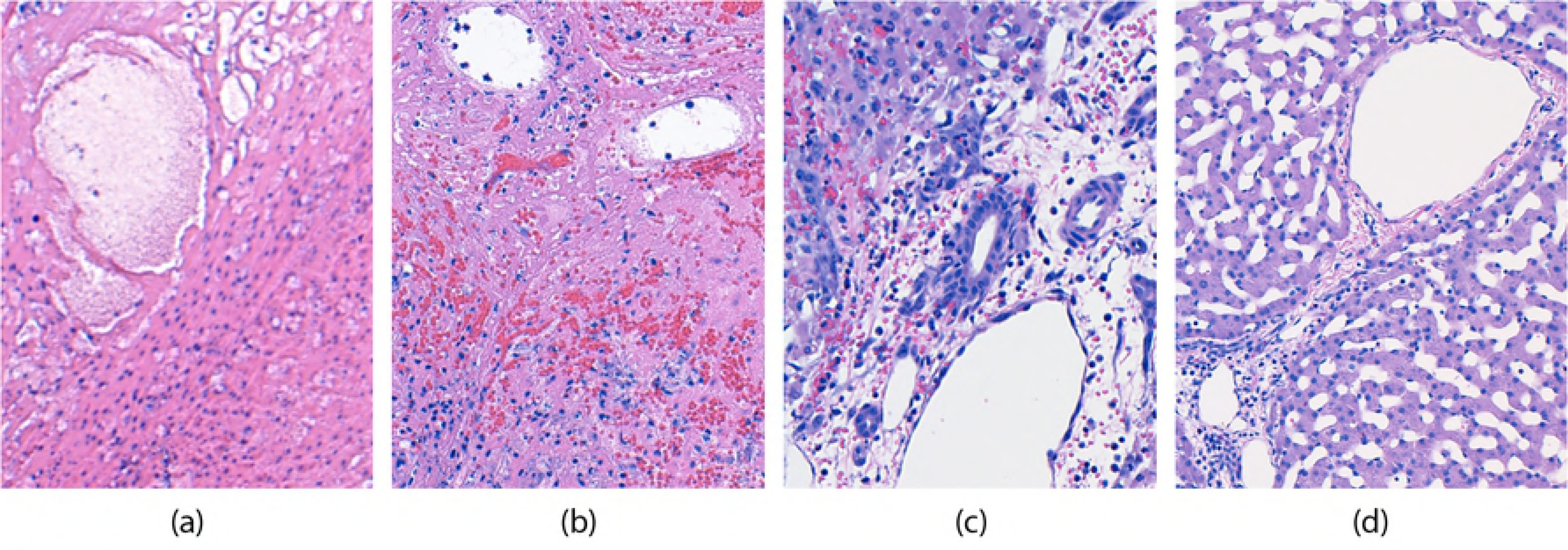
Histological results of the different gradients of necrosis and tissue damage as viewed histologically. (a) Histology image from Region A, Complete necrosis; (b) Region B, Complete necrosis with traces of hemorrhage; (c) Region C, Complete necrosis of hepatocytes but bile ducts in tact; (d) Region D, healthy liver tissue.

The minimum diameter of the necrotic region, from Region A out to Region C, was measured to be 2.08 cm as seen in Fig 8. As the plane of the tissue sections was not strictly perpendicular to the tract of the cryoprobe, the minimum diameter was used to determine the minimum cross sectional area of 3.4 cm^2^. As the iceball forms in a cylindrically symmetrical shape, any cross section will result in an ellipse with a minor diameter equal to the diameter of the cylinder.

## Discussion

The results of this study demonstrate the capacity of CO_2_-based cryoablation with a percutaneous cryoprobe to cause necrosis in large volumes of tissue and tumors. In tests performed in tissue phantom, a confidant freezing volume with a radius of 2.23 ± 0.075 cm was observed over six trials. A minimum temperature below −40 °C was achieved in all trials, which satisfies the standard freezing temperatures for tumor necrosis accepted for similar cryoablation devices currently used in the United States [16]. While the freezing capacity of the cryoablation device was verified in phantoms, *in vivo* studies were performed to determine whether the device could induce tumor necrosis.

Histopathology of carcinogen-induced mammary tumors in rats demonstrated the device’s ability to repeatedly induce necrosis in large volume malignancies. However, in the rat tumor model, there was not significant heat load. Thus, we evaluated the device’s efficacy in a swine liver, i.e., 39 °C - an environment with a heat load similar to that of the human breast, i.e., 37 °C [25]. This step was crucial in validating freezing ability because the heating effect of blood perfusion could significantly reduce iceball formation. Liver sections and histology confirmed that our CO_2_-based cryoablation device is capable of inducing necrosis in up to 2.08 cm diameter regions. Therefore, the performance of our custom built cryosystem is comparable to benchmark values of various accepted cryoablation standards of other gas-based systems in the United States.

While promising, there are several limitations to this study that will direct future work. Although results have shown cellular necrosis of a 1.5-cm diameter mammary tumor both in rats and in swine liver, it should also be noted that this study did not investigate the use of this device in the necrosis of cancerous tissue under a heat load. These preliminary studies did provide information related to the potential for damage to healthy tissue and skin, which could lead to issues in wound healing. This must be examined further in order to ensure the named device can match the standards of care set by existing cryoablation systems.

Future studies are warranted to examine several areas including the extent of necrosis in cancer under a heat load and the impact of the device on healthy tissue and long-term healing. Additionally, there have recently been significant inquiries into the immune response to breast cancers due to cryoablation. These include animal studies examining rejection rates of tumors in mice when rechallenged [26,27] as well as human trials examining the immune response and overall survival outcomes of patients treated with cryoablation [28,29]. This promising work furthers the potential for impact that cryoablation of breast cancer could have in LMICs, and opens the door for future studies to examine the immune response induced by this device.

## Conclusions

This study evaluated the use of a cryoablation device that utilizes CO_2_ as the cryogen in a variety of experimental settings, thus exploring its potential as a breast cancer treatment in LMICs. Although previous work has examined the use of argon-based cryoablation, no such work has explored the efficacy of a CO_2_-based cryoablation device. Our results demonstrate the efficacy of this device in phantoms, at the cellular level as examined in induced mammary tumors in rats, and under high heat load and perfusion as performed in swine liver. Thus, this device has passed crucial benchmarks meriting further research in both large animals with tumors and eventually in human trials. With the introduction of this device, we are optimistic that the care paradigm in LMICs may shift in the near future to account for the availability of affordable and effective treatment in rural areas.

## Acknowledgements

We would like to thank the Center for Bioengineering Innovation and Design and the Biomedical Engineering Department at Johns Hopkins University for research support, as well as: Monica Rex, Yechan Kang, Nikhil Jois, Alwin Hui, Sonia Trakru, Sarah Lee, Sanjay Elangovan, Ben Lee, Marton Varady, and Sue Lucas.

